# Extremely fine-scale soil heterogeneity in a rare serpentine endemic plant shapes patterns of genetic diversity

**DOI:** 10.1101/2025.09.01.673272

**Authors:** Joseph Braasch, Julia G. Harenčár, Sarah Swope

**Affiliations:** Department of Ecology and Evolutionary Biology, University of Arizona, Tucson, AZ 85721; Center for Population Biology, University of California, Davis, CA 95616; Mills College, Northeastern University, Oakland, CA 94613

**Keywords:** Landscape Genetics, Plant Ecology, Serpentine Ecology, Calochortus, Environmental Heterogeneity

## Abstract

Studies of population genetic structure are typically conducted at the scale of species distributions and encompass large distances and substantial environmental variation. However, population genetic structure could also be present in species with highly restricted global distributions, such as habitat specialists with threatened or vulnerable conservation status. For these organisms, low dispersal distances coupled with fine-scale environmental heterogeneity could influence population genetic composition, potentially creating spatial genetic structure and genotype by environment associations. Here we use the serpentine endemic plant *Calochortus tiburonensis*, with a global distribution of 160 ha, to evaluate whether fine-scale structure in soil composition and low seed dispersal distances result in the development of population genetic structure. We paired soil elemental analysis with a RAD-seq SNP dataset for 24 C. tiburonensis individuals. Although no population structure was detected between *C. tiburonensis* sampling locations, multiple analyses identified associations between soil composition and genetic distance between individuals. This included associations with nickel and magnesium, two elements that were expected *a priori* to impact plant fitness in serpentine landscapes. However, redundancy analyses and a generalized dissimilarity model both suggest that total soil variation better explains differences in genetic composition between individuals, implying that selection from the holistic soil environment has a role in matching plant genotypes to the microenvironment. Our results indicate that fine-scale environmental heterogeneity could influence genetic differences between individuals in plant populations, even in the absence of population genetic structure. Additionally, these associations between genetic composition and fine-scale environmental heterogeneity implicate extremely fine-scale environmental heterogeneity as an essential mechanism for preserving genetic variation, particularly within range-limited species.

## Introduction

Environmental heterogeneity contributes to the generation of biological diversity and population genetic structure (Hendrick et al. 1976, Smith et al. 1997, Ortego et al. 2012, Daleo et al. 2023). The full characterization of genetic structure requires consideration of the organism’s entire range and will naturally include a large spatial area for most species (Eckert et al. 2010, Freeland et al. 2010, Ortego et al. 2012, Anderson et al. 2010). However, environmental heterogeneity exists across a range of spatial distances and can contribute to genetic structure at fine scales, particularly when selection associated with heterogeneity is strong (Anderson et al. 2010). However, the term ‘fine-scale’ in this context can also refer to vastly different spatial distances and often includes survey distances greater than 50km (Gugger et al. 2019). Additionally, environmental heterogeneity can contribute to spatially variable selection, with or without the presence of spatial genetic structure (Hendrick et al. 1976, Freeland et al. 2010, Tigano and Friesen 2016, Hoey and Pinsky 2018). Yet, the importance of fine-scale heterogeneity on the scale of meters, and its relationship with genetic diversity and population genetic composition, is comparatively less understood and rarely investigated (Whitlock 2015) (but see Hens et al. 2013).

Serpentine soils are notable for their high within- and between-patch variation in soil composition (Krukeburg 1950). These soils often exist as a heterogeneous matrix embedded within landscapes derived from non-serpentine parent material. Additionally, serpentine soils are notable for possessing low quantities of essential nutrients (N, P, K, Ca), high quantities of heavy metals (Ni, Cr, Co), and having low water holding capacity, making them challenging and potentially toxic substrates for plant growth (Walker 1954, Proctor and Woodell 1975). These characteristics act as strong selective agents that many plants must evolve to tolerate (Anacker 2014, Cacho and Strauss 2014). Indeed, serpentine plant communities are commonly used to study plant adaptation to stressful soil environments (Krukeburg 1951, Wright et al. 2006), and unique alleles that confer resistance to heavy metal toxicity have been repeatedly identified in serpentine plant populations (Niu et al. 2018, Celestini et al. 2025). Importantly, there is fine-scale heterogeneity in the composition of serpentine soils, with considerable turnover at the scale of meters (Krukeburg 1950). These large changes in the abundance of heavy metals are expected to impact plant fitness, producing a matrix of stress within serpentine outcrops that also varies based on plant genotype.

Serpentine communities support a high proportion of endemic plant species (Krukeburg 1984, Skinner and Pavlik 1994, Safford et al. 2005), which now face existential threats from anthropogenic climate change (Harrison et al. 2015). In California, the stressful nature of these soils appears to have buffered plant communities from the invasion of Eurasian grasses and forbs that have transformed the majority of grassland communities (Huenneke et al. 1990, Harrison and Inouye 2002, Burge et al. 2016). Consequently, serpentine plants are not only useful for the study of plant adaptation and stress; they are also a conservation priority. Many serpentine endemic species do not maintain viable populations on non-serpentine substrates, which coupled with low dispersal distances, could inhibit expansion or range shifts in these species and leaves adaptation as the primary tool through which serpentine endemic plants can persist in a warming world (Jump and Peñuelas 2005). Thus, knowledge of how environmental factors, such as soil heterogeneity, influence the genetic composition of serpentine endemics could be crucial to their conservation.

*Calochortus* is an iconic wildflower genus that underwent an adaptive radiation across the California Floristic Province and includes many serpentine specialists with restricted ranges. *Calochortus tiburonensis* has one of the smallest estimated natural geographic distributions of any plant, with a range restricted entirely to a single coastal hilltop – the Ring Mt. Preserve. This preserve protects 160 ha, of which approximately half is serpentine grassland, while the other half is residuum weathered from sandstone and shale, supporting both grassland and woodland communities. In addition, *C. tiburonensis* is found only on north- and east-facing slopes. Hence, the persistence of the species hinges on this singular population with approximately 18 ha of potential habitat. Conservation in situ is therefore crucially important, because A) the genetic diversity present in this population must be sufficient for any necessary adaptation to future conditions, and B) the species cannot migrate to other sites, nor can it be restored with propagule from outside locations. Thus, understanding the relationships between genetic diversity and environmental variation can be used to improve management outcomes for the singular *C. tiburonensis* population.

We use RAD sequencing to produce a SNP dataset assembled from C. tiburonensis individuals across the species’ entire 1.6 km^2^ global distribution. Using this dataset, we first test whether distinct *C. tiburonensis* patches correspond to genetic structure caused by low seed dispersal distances. Second, we ask whether the genetic composition of *C. tiburonensis* individuals corresponds to components of the soil environment. To test this hypothesis, we compared the genetic composition of *C. tiburonensis* individuals within a multivariate framework in addition to testing for the explanatory power of each soil component on the genetic differences between individuals. We expected that metals previously shown to be associated with plant fitness in studies of serpentine adaptation, such as magnesium and nickel, to have the greatest power in predicting *C. tiburonensis* genetic composition. Our study will help quantify the importance of small-scale environmental variation in determining plant genetic variation. During periods of rapid anthropogenic environmental change, a more complete understanding of the processes that structure genetic diversity will be integral to the conservation of *C. tiburonensis* and many other organisms.

## Methods

### Study system

*Calochortus tiburonensis* A.J. Hill (Liliaceae) is a long-lived perennial plant that grows from a bulb and produces a single, long, strap-like leaf and a single scape that typically produces 1 – 5 flowers (range 1 – 12; Swope, *unpublished data*) (Figure 1). In any year, mature plants may flower, or they may be vegetative (produce a single leaf but no flowers), or they may be dormant with no aboveground tissue. Plants often enter dormancy for one or more years after flowering (Swope, *in preparation*). Flowers are large, showy, and visited by generalist insects (Bombus species, Apis mellifera, and species of Syrphidae and Coleoptera). The effectiveness of these visitors as pollinators is unknown. Reproduction from seed is episodic; plants are also capable of cloning via bulb splitting, although the frequency of this is low, especially relative to germination from seed (Swope, *unpublished data*). Seeds are heavy and have no obvious adaptation to promote long-distance dispersal either by wind or by animals.

**Figure 1.**
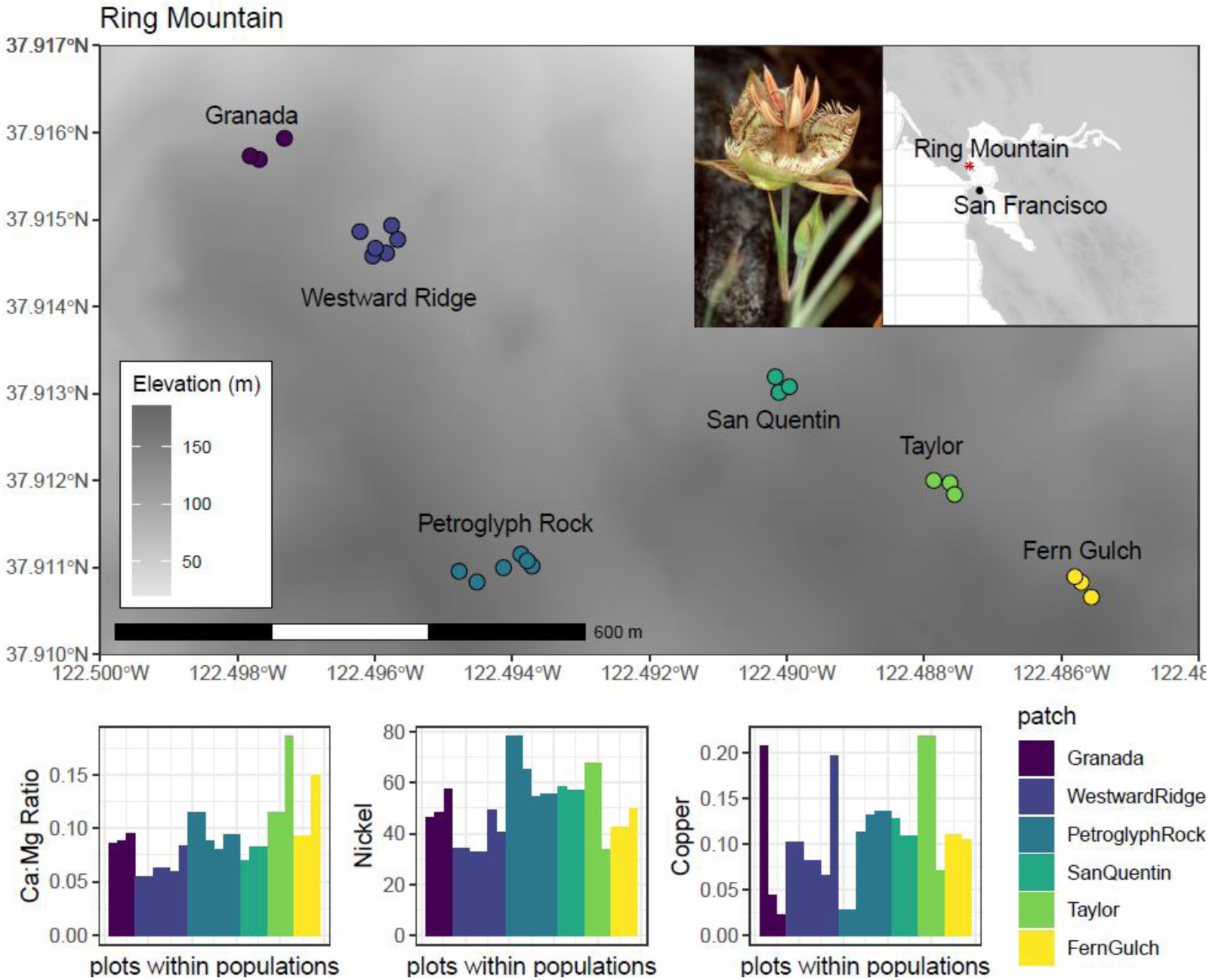
Top: Map showing the sampling locations of *Calochortus tiburonensis* across Ring Mountain in California, USA. Names are assigned to six different patches of *C. tiburonensis* where soil and plant material were collected. The number of soil samples collected from each patch depended on patch size, ranging from 3-6 samples per patch. Points within the map are jittered to better indicate the existence of plots in close proximity. Bottom: Barplots demonstrating variation in soil composition at small spatial scales for potentially phytoxic elements. Individual bars indicate the amount of each chemical across all 24 sampling locations. Plots are colored by patch in all figures. Photo credit for *C. tiburonensis* image: Rick York and the California Native Plant Society.

*Calochortus tiburonensis*’ entire geographic range is a single hilltop known as Ring Mountain (Marin County, CA, USA). This highly restricted geographic range appears to be natural, and not the result of anthropogenic habitat loss, given the extensive activity of foundational botanists in California and no reports of its presence elsewhere. *C. tiburonensis* grows exclusively on serpentine outcrops, which are patchily distributed and comprise ∼18 ha of the ∼50 ha Ring Mt. Preserve. It is listed as Threatened at both the State and Federal levels.

### Soil samples and chemistry

We collected a single soil sample associated with each of 24 long-term demography plots randomly located within the six distinct patches of *C. tiburonensis* present on Ring Mt. Preserve (Figure 1). While there is one additional small patch, it was not sampled due to its proximity to a well-used hiking trail. Plots within individual demography plots were often separated by ∼1-5m to capture the fine-scale resolution of soil heterogeneity. Soil samples were collected using a 5 cm × 15 cm cylinder attached to a slide hammer. Each sample was placed and stored in its own bag. Soils were then air-dried at ambient temperature, sifted to remove rock fragments, and analyzed for 24 essential nutrients and heavy metals at the Cornell Soil Health Laboratory (Cornell, New York, United States).

Assessments of how individual soil components contribute to genetic patterns could be confounded by correlations in the abundance of different soil elements. To identify which soil chemicals were correlated with one another, we calculated the Pearson’s correlation coefficients and plotted correlations with the *pairs.panels* function from the psych package in R. Because this generates a large matrix that impedes interpretation, we subset the data to only include chemicals frequently associated with plant fitness (magnesium, calcium, and nickel), chemicals that showed up as potentially important in further analyses (below; copper and silicon), and chemicals correlated with any of those previous chemicals with a Pearson’s correlation coefficient greater than 0.7.

### Plant Tissue Sampling

We collected several centimeters of tissue from the fresh green leaf of 24 randomly selected plants growing nearest to each soil sample. Two plant tissue samples from Petroglyph Rock (Figure 1) were collected directly next to each other and associated with the same soil sample. This was intentionally done to identify if adjacent plants are expected to be clones produced via bulb splitting. Leaf tissue was stored in separate coin envelopes, in the dark, at room temperature with silica beads (PolyLamProducts, Williamsville, NY) and extracted one to two weeks after collection. Genomic DNA was ground by mortar and pestle and extracted using Qiagen’s DNeasy Plant kit (https://www.qiagen.com)(Qiagen, Hilden, Germany) according to manufacturer instructions. The tissue collected from a single individual was divided and extracted in multiple rounds, then cleaned with an ethanol precipitation and pooled to generate sufficient high-quality DNA for sequencing. RAD sequencing and library preparation were conducted by Floragenex (Eugene, Oregon).

### De Novo Assembly and SNP calling

Floragenex fragmented the genomic DNA using the restriction enzyme PstI and individually barcoded each individual. The reduced representation library was then sequenced using an Illumina HiSeq platform (Illumina, Inc., San Diego, CA, USA). Custom scripts were used to de-multiplex raw reads, which were then cleaned with the package SNOWHITE 2.0.2 (Dlugosch et al. 2013).

*De novo* locus assembly and SNP calling were performed with STACKS 1.20 (Catchen et al. 2011; Hohenlohe et al. 2011) using the *denovo_map.pl* pipeline. Locus assembly was performed requiring a minimum of five reads (m = 5), a minimum of two mismatches (M = 2) consistent with expectations for diploid biallelic loci, and a minimum of two polymorphisms per individual (n = 2). The populations.pl module was then used to filter the set of SNPs requiring each locus to be present in at least three individuals per putative population (p = 3) and the assembled stack in at least 40% (two) of the six putative populations. When multiple SNPs were identified within a single assembled locus, we randomly selected a single SNP to include in our final dataset using the option ‘write_random_snp’. The populations.pl module was also used to calculate population level summary statistics: observed heterozygosity (H_o_), inbreeding coefficient (F_is_), and nucleotide diversity (π).

### Population structure and genetic differentiation

The apparent isolation of *C. tiburonensis* patches could result in genetic clustering if seed and pollen dispersal are low, despite the limited 1.6 km2 range of the species. We used the Bayesian clustering program STRUCTURE v 2.3.3 (Pritchard et al. 2000) to identify the presence of distinct genomic groups across the 24 sampled individuals. We tested for k distinct subpopulations ranging from one to seven, with seven being one greater than the number of sampled patches. The admixture model was chosen for this analysis, which allows for no *a priori* population assignment. Each value of k was simulated over ten independent runs for 100,000 generations after a 10,000 generation burn-in period. We used CLUMMP v.1.1.2 (Jakobsson and Rosenberg 2007) to compare the likelihood estimate for each value of k by averaging across model runs. The most likely value of k is determined by the set of runs with the highest overall similarity, summarized as H’.

Spatial clustering could also be apparent in multivariate analyses and visualizations of genetic diversity. To assess genotypic differences across all sampled individuals, we used principal component analysis (PCA) to identify the number of genetic clusters present across multivariate axes of allele frequencies. We used the function ‘find.clusters’ in the R package *adegenet* (Jombart and Ahmed 2011), which uses unsupervised machine learning to minimize the distance between individuals and potential population centers within a PCA. The maximum value of k was again set to seven for this analysis. We also visualized genetic clustering of *C. tiburonensis* individuals in PCA space with the function ‘dudi.pca’. Multivariate methods, including PCA, often require data with no missing observations.

However, missing allele calls are common in RAD-seq data sets. Missing allelic assignments in the dataset were imputed by replacement with the most common allele across all individuals for each locus.

### Associations between genomic composition, distance, and soil chemistry

Clustering analyses identified only one genetic cluster that included all sampling sites (see Results). We therefore conducted all subsequent analyses assuming a single population. Additionally, because two *C. tiburonensis* individuals from the Petroglyph Rock shared a paired soil sample, one individual was randomly removed prior to conducting any analyses including soil composition.

Testing for associations between the genomic composition of plants and the abundance of different soil compounds was performed using three approaches; Mantel tests, distance based redundancy analysis (dbRDA), and a generalized dissimilarity model (GDM). Each test uses distinct and complementary approaches to identify associations between the genetic distance matrix and matrices of explanatory variables (Legendre & Anderson 1999, Fitzpatrick & Keller 2015). Mantel tests assess the strength and significance of the correlation between two matrices and can be used to compare genetic distances to the differences in the abundance of individual soil elements. These tests were used to identify correlations between the matrix of genetic distances between individual *C. tiburonensis* and the difference in single components of soil variation. Each test was conducted independently using the mantel function in base R, with 999 permutations and assessed by the Pearson correlation coefficient. The second approach, dbRDA, is a constrained ordination that tests for linear relationships between predictor variables (soil composition) and a multivariate dataset (genetic composition). Importantly, in addition to conducting the analysis along the PC axis of genetic variation that is most strongly correlated with the predictor variables, it is possible to include multiple soil variables simultaneously in the analysis. In comparison, the third approach, GDM, tests for non-linear associations between the change in predictor variables and the turnover in genomic composition. We expect that soil compounds associated with genetic differences in the Mantel tests and dbRDA will also be important contributors to the GDM model. However, soil compounds that have non-linear relationships with *C. tiburonensis* genetic composition might only appear using GDMs.

To implement dbRDA, we used the matrix of alleles with imputed assignments for null calls (36.4% of all cells in the matrix of genetic identity across individuals). This is because, similar to PCA, RDA cannot be performed with missing data. Although there were many soil compounds that could be included in this analysis, RDA models can be overfit and should only be used with uncorrelated variables (Capblanq & Forester 2021). We therefore chose to run two models with a reduced number of predictor variables: 1) a model with Magnesium, Calcium, and Nickel and 2) a model that used the first three principal component axes of soil variation as predictors. The first model allows us to test our a priori expectations for elements that are commonly known to be correlated with plant fitness and plant adaptation to serpentine environments. The second model allows us to consider the entire soil environment and can identify when variation in total soil composition is correlated with genetic differences, in addition to suggesting which compounds contribute most to this relationship. However, this framework does not allow for the identification of which compounds, individually or synergistically, drive the relationship. Both dbRDA models were performed using the function ‘dbrda’ from the R package *vegan* (Oksanen et al. 2022) using Euclidean distances. To test whether soil variation was significantly associated with RDA axes, we performed an ANOVA for each model with 999 permutations.

Soil environments are complex and composed of many elements and compounds. The framework of a generalized dissimilarity model allows for the testing of many explanatory variables simultaneously and, importantly, can identify the presence of non-linear relationships between predictor variables and the response matrix. This capacity is modeled via the i-spline functions produced with the model, showing the linear, exponential, logarithmic, or cubic functions that describe the relationship between turnover in the response matrix and the ecological gradient. It is the maximum y-axis value of these functions that also determines the relative contribution of each variable towards describing genetic differences. This modeling approach also incorporates the spatial distance between samples to test for isolation by distance. To produce a distance matrix to serve as the response variable in the GDM, we calculated Euclidean distances using the function ‘vegedist’ from the *vegan* package. To test for model significance, we used matrix permutation to compare the distribution of deviance explained using permuted data (500 permutations) to the deviance explained in the un-permuted dataset. The relative importance of each soil component was evaluated as the maximum y-axis value for the corresponding i-spline function. We implemented the GDM and related analyses using the package *gdm* (Fitzpatrick et al. 2022).

## Results

### Soil Composition

Pairwise comparisons of select soil components identified seven significantly correlated elements (Figure 2). Surprisingly, six of these correlations all involved copper, which was positively correlated with calcium, magnesium, and nickel which are all known to impact serpentine plant ecology. Additionally, copper was also positively correlated with boron and aluminum and negatively correlated with strontium and arsenic. Lastly, nickel was also positively correlated with aluminum.

**Figure 2:**
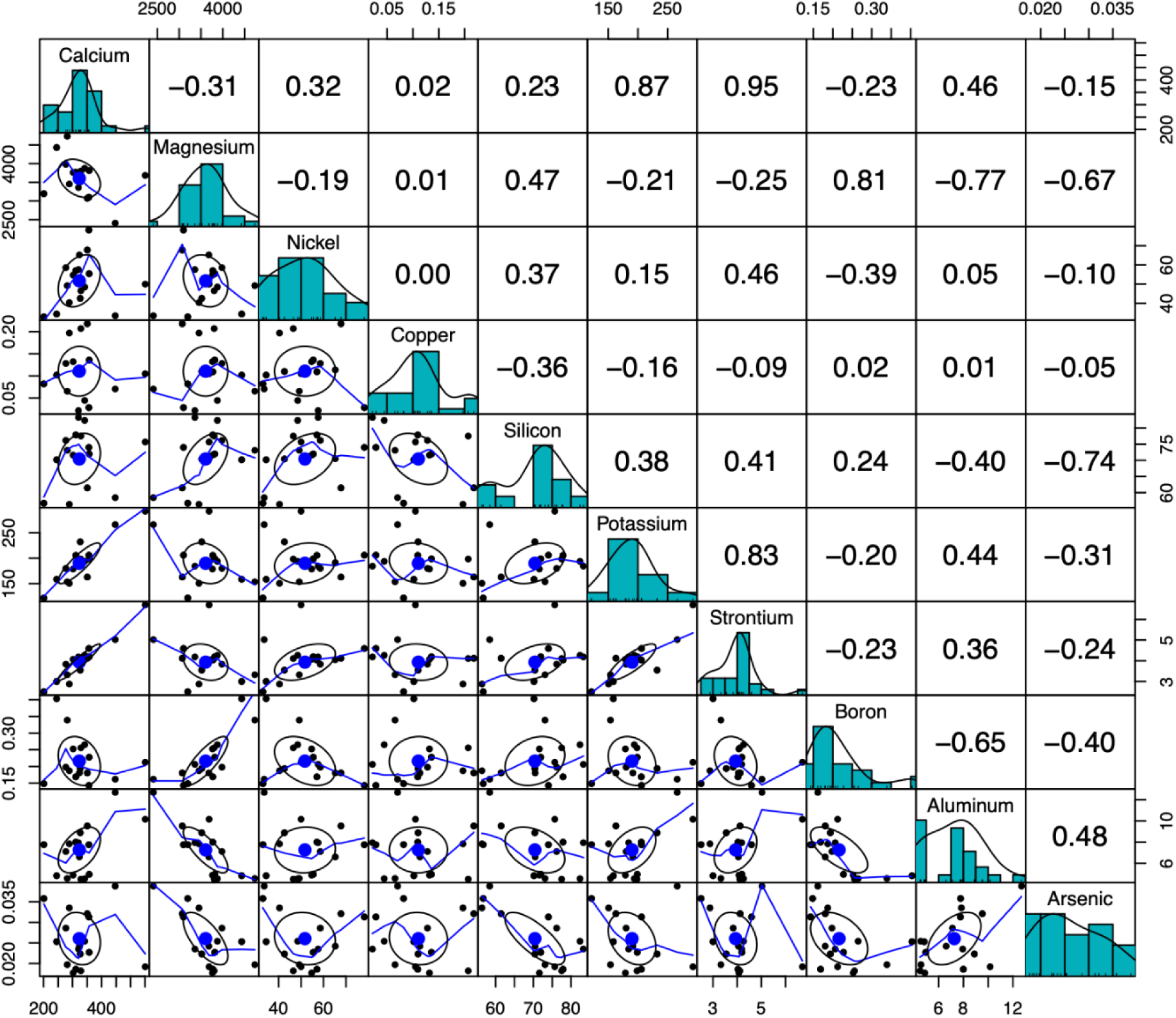
Correlations in the abundance of soil chemicals, subset to highlight elements that are expected to affect serpentine plant fitness (Calcium, Magnesium, Nickel, and Copper) and elements they are significantly correlated with. Bivariate scatter plots with loess smooth lines and correlation ellipses are displayed below the diagonal, histograms of soil value distributions are on the diagonal, and the Pearson correlation value is above the diagonal.

### Assembly and genetic diversity

STACKs identified 128405 biallelic loci across all 24 individuals. The reduced dataset, which contained only loci with allele calls for all individuals, contained 452 loci. The genetic diversity of *C. tiburonensis* sampling sites was greatest in sampling sites with greater population densities and higher sampling effort (Table 1). These sites also possessed the highest inbreeding coefficients, consistent with these sites possessing higher proportions of individuals that are homozygous at the sequenced sites.

**Table 1.** Estimates of genetic diversity across *Calochortus tiburonensis* sampling sites, calculated across all nucleotide sites and only polymorphic sites assembled in STACKs.

|  | Site Name | n | H<sub>o</sub> | H<sub>e</sub> | F<sub>is</sub> | Pi | % Polymorphic Sites |
| --- | --- | --- | --- | --- | --- | --- | --- |
| All nucleotide sites | Granada | 3 | 0.0008 | 0.0013 | 0.0014 | 0.0016 | 0.3099 |
|  | Westward Ridge | 6 | 0.0008 | 0.0016 | 0.0021 | 0.0018 | 0.4539 |
|  | Petroglyph Rock | 6 | 0.0008 | 0.0015 | 0.0021 | 0.0017 | 0.4365 |
|  | San Quentin | 3 | 0.0006 | 0.0011 | 0.0013 | 0.0014 | 0.2555 |
|  | Taylor | 3 | 0.0008 | 0.0012 | 0.0013 | 0.0016 | 0.3072 |
|  | Fern Gulch | 3 | 0.004 | 0.0010 | 0.0014 | 0.0012 | 0.3072 |
| Polymorphic sites | Granada | 3 | 0.1023 | 0.1647 | 0.1766 | 0.2075 |  |
|  | Westward Ridge | 6 | 0.1047 | 0.2023 | 0.2722 | 0.2282 |  |
|  | Petroglyph Rock | 6 | 0.0993 | 0.1978 | 0.2702 | 0.2250 |  |
|  | San Quentin | 3 | 0.0745 | 0.1405 | 0.1714 | 0.1806 |  |
|  | Taylor | 3 | 0.1071 | 0.1628 | 0.1640 | 0.2039 |  |
|  | Fern Gulch | 3 | 0.0484 | 0.1223 | 0.1751 | 0.1583 |  |

### Population structure and spatial clustering

There was no evidence of clonal reproduction across the individuals sampled.

Bayesian clustering analysis implemented in STRUCTURE assigned similar likelihoods for the existence of one and two populations (H’=0.393, H’=0.396 for K=1 and K=2 respectively). Although K=2 had slightly greater support, it predicted an entirely admixed population from two unsampled subpopulations, which is biologically implausible, because our sampling encompassed the entire distribution of *C. tiburonensis*. The likelihood values for K=1 and K=2 were similar, with Bayesian criterion information scores of 231 and 233 for one and two populations, respectively. Again, the population assignments for K=2 were dubious, with one individual being assigned to a second population while all other individuals shared assignment to the first population.

We obtained similar results from a PCA based analysis of structure (Figure 3). Although PCA of SNP data often depicts only a small proportion of the total variation, a consequence of the incredibly high dimensionality of the data, it is still useful for visualizing the genetic similarity of individuals. In our dataset, the first four PC axes explained 6.3, 6.1 5.8, and 5.8% of the total variation, respectively. Within this framework, we found that most individuals clustered around the origin of the PCA plots, with one or two individuals distinctly separated along each axis (Figure 3). This behavior is indicative of low genetic differentiation because the largest PC axes do not depict any distinct clusters, as indicated by prior clustering analyses.

**Figure 3.**
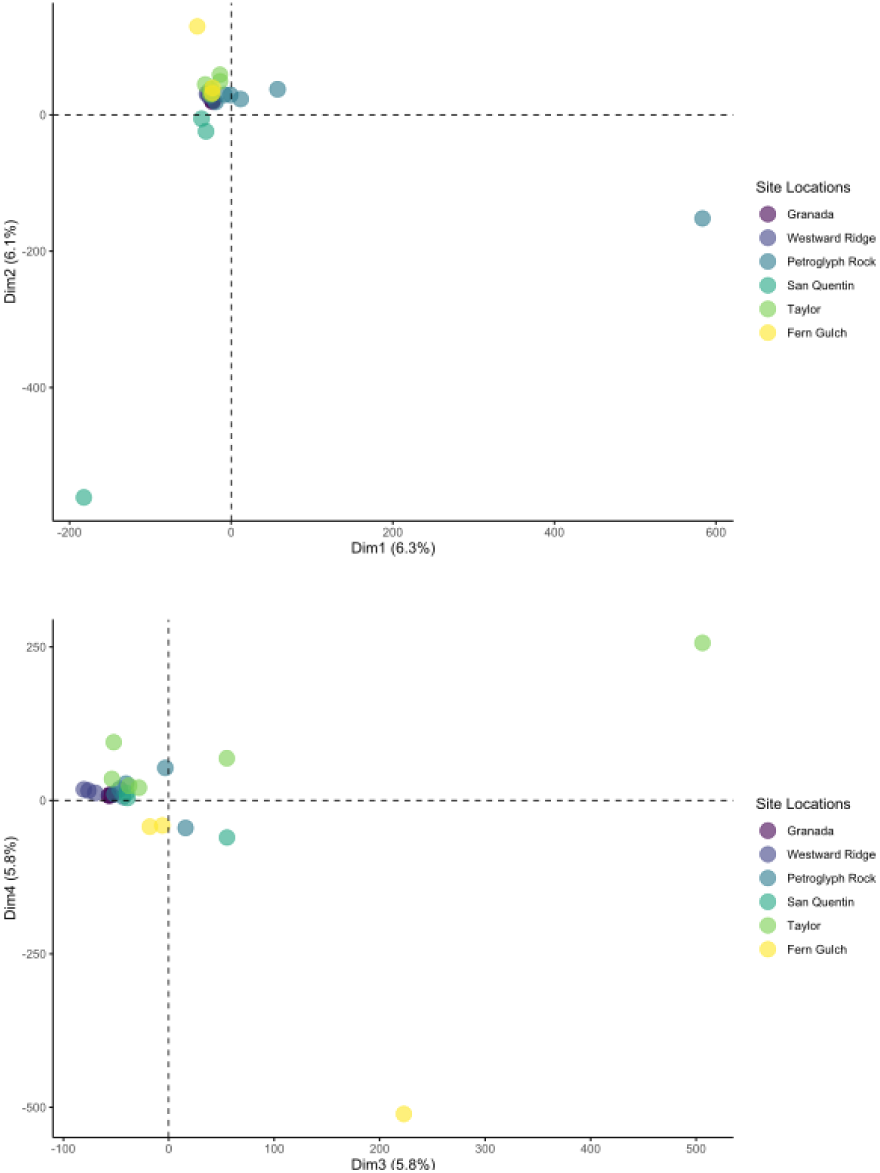
Principal components analysis of *C. tiburonensis* SNPS. The first four PC axes explained similar amounts of the total genetic diversity and explained 24% of total variation.

### Associations between genomic composition, distance, and soil chemistry

Of the 28 components of soil composition variation, Mantel tests identified four that were significantly correlated with differences in genetic composition among individual *C. tiburonensis* (Table 2). Three of these soil elements were a priori expected to correspond with plant fitness; copper, magnesium, and nickel. The fourth correlate, silicon, was not predicted to correspond to plant fitness.

**Table 2.**
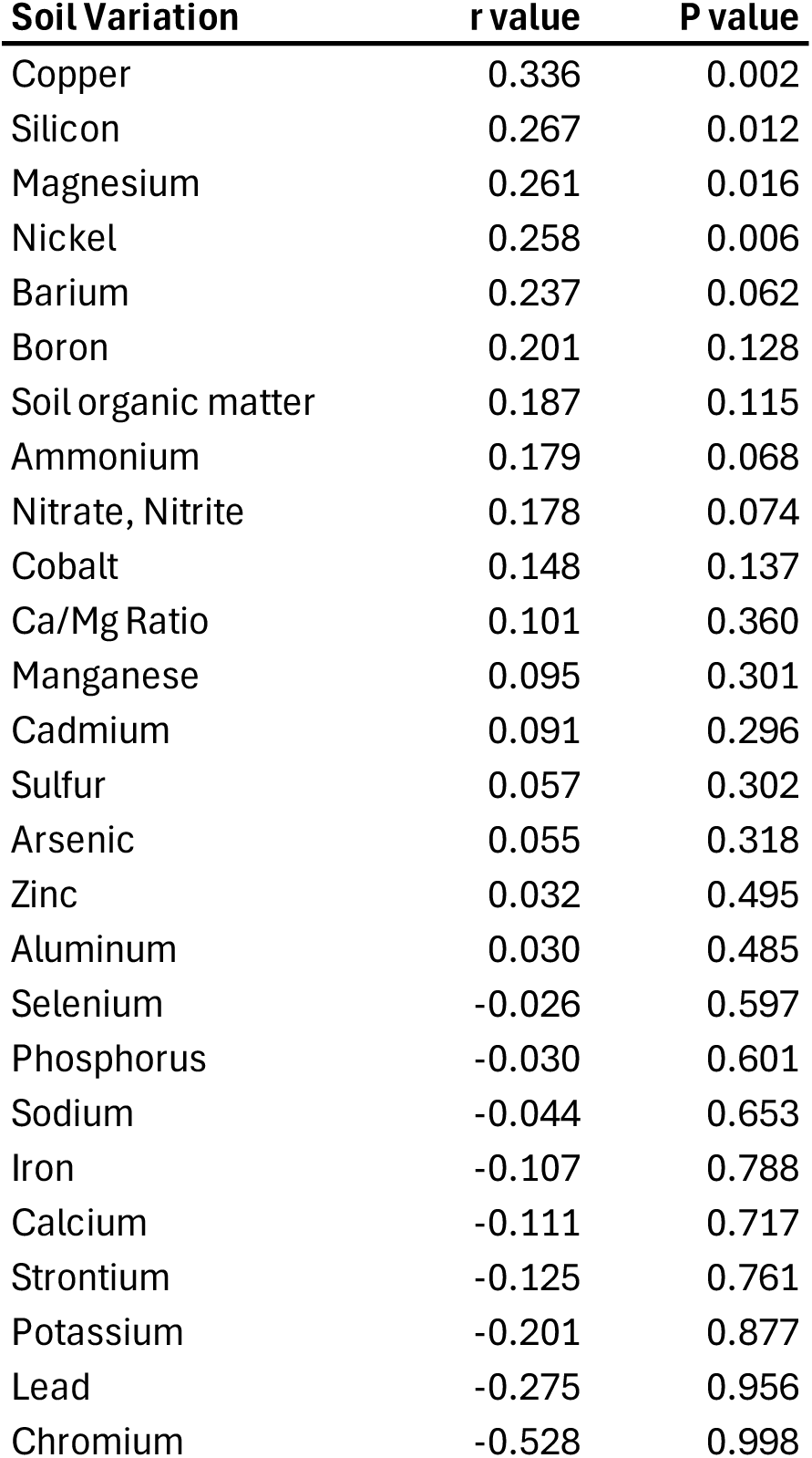
Mantel results testing for the correlation between genetic distances and differences in soil composition, ordered from highest to lowest correlation coefficients.

We used two different dbRDA models to test for correlations between the genetic composition of *C. tiburonensis* individuals and variation in soil chemistry. The first model included the abundance of magnesium, calcium, and nickel, three soil elements commonly associated with plant fitness on serpentine soils. The full model was not significant (F = 1.07, P = 0.13), although the first RDA axis, which was most strongly associated with soil magnesium content, explained a significant proportion of variation (Table 3, Figure 4A). The second dbRDA model was built using the first three PC axes of variation in soil composition, which explained 72.6% of the total soil variation (Figure 4B). This model, which incorporated all variation in soil composition, was significant (F = 1.11, P = 0.009). Additionally, the first RDA axis explained a significant proportion of genomic variation when ordination was constrained by the additive model, while the second and third RDA axes did not (Table 3, Figure 4B).

**Figure 4.**
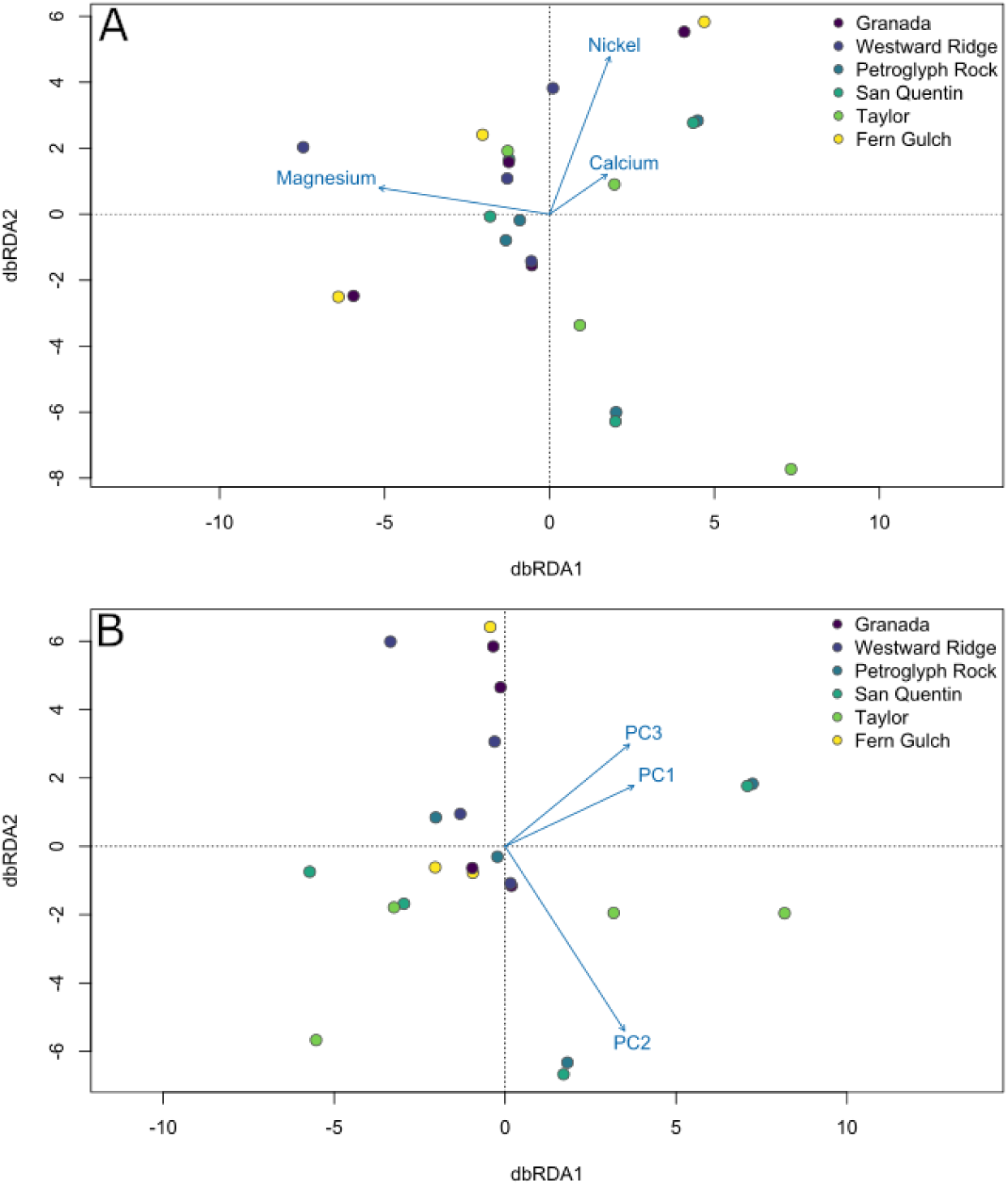
Redundancy Analyses of *C. tiburonensis* genetic diversity modeled with metrics of soil composition. A) An RDA modeled using three elements, Mg, Ni, and Ca, known to influence plant fitness on serpentine soils. The complete RDA model is not significant, although the effect of Mg is. B) An RDA modeled using the first three principal components of total soil variation. The full model is significant in addition to the first PC axis.

**Table 3.** Model results for two redundancy analysis using additive models of soil variation to constrain multivariate genomic variance. The first model used three soil elements commonly associated with serpentine plant fitness. The second model used the first three principal component axes of soil variation.

| Model | RDA Axis | Variance Explained | F value | P value |
| --- | --- | --- | --- | --- |
| Mg+Ca+Ni | Axis 1 | 0.38 | 1.21 | 0.006 |
|  | Axis 2 | 0.34 | 1.09 | 0.573 |
|  | Axis 3 | 0.27 | 0.89 | 0.781 |
| PC1+PC2+PC3 | Axis 1 | 0.37 | 1.20 | 0.016 |
|  | Axis 2 | 0.33 | 1.12 | 0.349 |
|  | Axis 3 | 0.30 | 1.00 | 0.540 |

A GDM that used turnover in soil composition to predict turnover in genetic composition found that soil explained a significant proportion of GDM deviance (41.8% deviance explained, p = 0.002). A set of 11 soil components contributed to the GDM model (Figure 5A). These components include magnesium and nickel, in line with our expectations for which serpentine soil elements should impact plant fitness. Additionally, organic compounds and nitrates, which should correspond to the nutritive quality of the soil, were also included. Notably, spatial distance and calcium were not assigned any predictive contribution to the final model. The former suggests no isolation by distance is present across the samples, which is unsurprising given the limited spatial distribution of *C. tiburonensis* and single population identified by cluster analyses. However, no individual soil component included in the model explained a significant proportion of the variance alone, and p values ranged from 0.30 to 0.822. This suggests that it is the combination of these soil components, and potentially their interactions, that are important for describing the genomic correlations with the soil environment.

**Figure 5.**
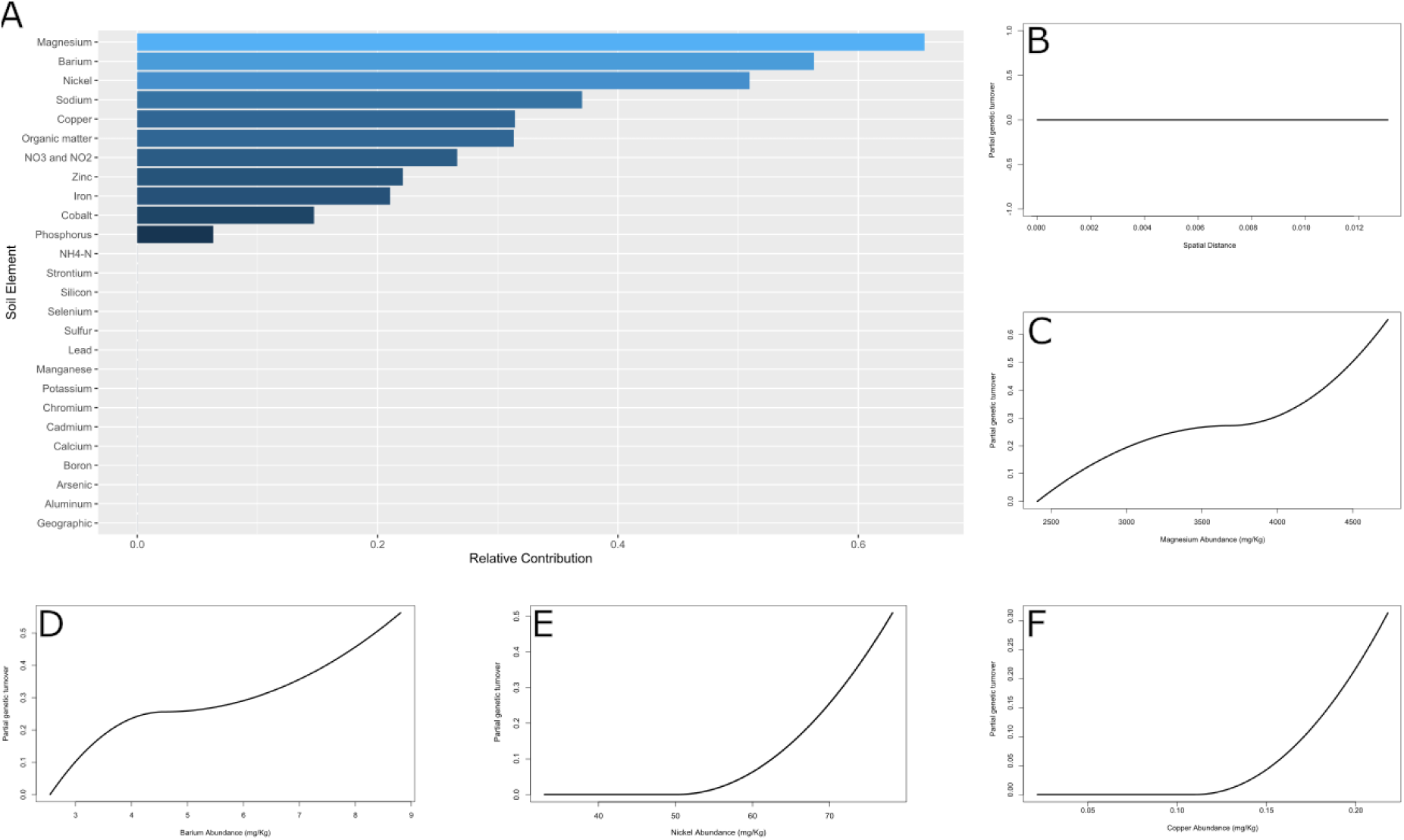
A generalized dissimilarity model identified 11 soil components as predictors of turnover in *C. tiburonensis* genetic composition. A) The relative contribution of all soil components towards explaining turnover in genetic composition. B) A flat isocline for genetic differences over increasing spatial distances is flat, further suggesting a lack of spatial structure. C-F) Isoclines depicting the non-linear relationships between increasing differences in the abundance of magnesium, barium, nickel, and copper.

## Discussion

Spatiotemporal heterogeneity has been shown to maintain genetic diversity associated with fitness across habitats and can contribute to the formation of spatial genetic structure (Freeland 2010, Ortego et al. 2012, Hens et al. 2013, Whitlock 2015). Our ddRAD-seq dataset of *C. tiburonensis*, a critically imperiled serpentine endemic herb growing on heterogeneous soils, identified no spatial genetic structure across the global distribution of the species. However, multiple lines of evidence suggest that soil composition influences genetic composition. Elements associated with plant stress on serpentine soils, such as magnesium and nickel, were correlated with *C. tiburonensis* genetic differences in univariate and multivariate tests. Copper, an essential element that can be toxic at high concentrations, was also found to be significantly correlated with genetic composition.

However, multivariate analyses suggest the total soil environment is a better predictor of *C. tiburonensis* genomic differences. In aggregate, our analyses suggest that fine scale soil heterogeneity drives variation in the genetic diversity of serpentine, and perhaps other, plant populations.

### Spatial Genetic Structure and Environmental Heterogeneity

The sole population of *C. tiburonensis* is restricted to the serpentine outcrops sampled in this study. The general absence of *C. tiburonensis* individuals outside of the sampled areas, in conjunction with exceedingly low seed dispersal distances, could have facilitated the formation of genetic neighborhoods. None of our analyses supported this hypothesis, suggesting that gene flow, likely via insect pollination, is sufficient to eliminate any spatial population genetic structure. Bees, the presumed primary pollinator of *C. tiburonensis*, have been found to transport pollen between flowers separated by over 1km (Jha and Dick 2010), which would easily allow for traversal across the entirety of *C. tiburonensis*’ global distribution. Indeed, analyses of genetic structure most strongly supported the presence of a single population, composed of all sampled areas, without meaningful differentiation across spatial distance. Consequently, any environmental factors that drive evolution within this species, are expected to be distributed without meaningful spatial structure across the landscape. This matches the expectations for serpentine soils, which are well known for their compositional heterogeneity and impacts on plant fitness.

Environmental heterogeneity can also generate spatial genetic structure, but few studies have tested for associations between genetic composition and the environment over small spatial scales as small as those investigated here. In one notable exception, differences in solar radiation across plots of the European Alpine plant *Biscutella laevigata* separated by ∼12m were significantly associated with genetic structure (Parisod and Christin 2008). However, environmental heterogeneity reinforced the existence of isolation by distance in *B. laevigata*, contributing to the presence of spatial genetic structure. In contrast, our analyses identified no spatial genetic signal despite significant correlations between soil and genetic composition. This outcome could be caused by the mosaic nature of environmental heterogeneity on Ring Mt. and the large turnover in soil composition at the scale of meters (Figure 1). A similar phenomenon has been observed in marine organisms, where larvae are widely and passively dispersed along ocean currents (Siegel et al. 2003), which can eliminate or reduce the presence of population structure, but adults segregate into different habitats based on their genotypic composition. These genome by environment associations have been observed in red abalone (*Haliotis rufescens*), marine snails (*Chlorostoma funebralis*), summer flounder (*Paralichthys dentatus*), and copper shark (*Carcharhinus brachyurus*) without the presence of spatial genetic structure (De Wit and Palumbi 2012, Gleason and Burton 2016, Hoey and Pinsky 2018, Klein et al. 2023). While similar effects might be expected in plants with large dispersal distances and high rates of gene flow, such as wind pollinated trees and grasses, these organisms appear to typically develop spatial genetic structure alongside environmentally driven genetic differences (Gugger et al. 2019). However, most studies also do not investigate spatial genetic structure over distances under 1km. Thus, the associations between soil and genetic composition in *C. tiburonensis* might not be an outlier in plant ecological genetics and might only be apparent due to the unique resolution of these data. Given that similar relationships between soil and genetic composition could exist across plant species, a more complete understanding of plant population genetics requires studies that investigate environmental heterogeneity over a large range of spatial resolutions (Manel et al. 2010, Gugger et al. 2019).

### Genetic Association with Soil Elemental Composition

Our analyses identified significant associations between potentially toxic soil elements, including magnesium, nickel, and copper, and genetic variation in *C. tiburonensis*. Heavy metals such as these have been well studied in serpentine plant ecology, where their abundance in the soil is toxic to many plants. However, papers reporting an effect of copper on serpentine plants are few, likely due to small differences in the abundance of copper between serpentine outcrops and nearby soils derived from other rock substrates (Wright et al. 2006, Oze et al. 2008). Soil copper abundance at Ring Mt. was similar or lower than at other well studied serpentine sites and also does not differ from quantities present at nearby non-serpentine sites (Wright et al. 2006, Oze et al. 2008, Lazarus et al. 2011). In comparison, the abundance of nickel and magnesium on Ring Mountain exceed the expectation for non-serptentine soils, although they were equivalent or lower than what has been measured in many other serpentine communities (Wright et al. 2006, Oze et al. 2008, Lazarus et al. 2011). This suggests that correlations between genetic composition and either magnesium or nickel are related to the toxic properties of elements on serpentine soils. However, associations between *C. tiburonensis* genetic composition and copper might not be a product of the serpentine environment. Since copper abundance is similar in nearby non-serpentine areas, similar relationships between copper and genetic composition could be present in nearby non-serpentine plant populations. The association of genetic composition with copper could also be due to strong correlations between copper and six other soil chemicals, including calcium, magnesium, and nickel (Figure 2).

Other elements, such as silicon and barium, were also associated with differences in genetic composition in one or more of our analyses. These elements are not expected to impact plant fitness and are not considered important for serpentine ecology. In addition, the first PC axis of soil variation was significantly associated with genetic composition. The first PC axis of soil variation was positively correlated with copper in addition to other elements that were not a priori expected to affect serpentine plant fitness, such as cadmium and sulfur. These soil elements could be ecologically relevant, but without effects that are unique to serpentine soils. For example, higher barium soil content has been observed in urbanized areas due to increased environmental contamination from manufactured materials and vehicle air pollutants (Monaci et al. 2000), consistent with Ring Mountain’s proximity to the city of San Francisco. However, an effect of barium on plant fitness has not been widely reported in the literature and barium soil variation might be similar in non-serpentine areas if it is of anthropogenic origin. In this regard, our results suggest that broadscale soil variation contributes to the genetic composition of plant populations. Genetic studies performed at the same scale to the study presented here would be required to confirm whether similar patterns are observed in other plant species, on and off serpentine soils.

Within the framework of a GDM, there were notable differences in the isocline shape for individual elements. Of the elements a priori predicted to be correlated with genetic composition, nickel and magnesium contributed the greatest predictive power, but had differently shaped isoclines. The nickel isocline was fit as an exponential curve, indicating that only large differences in the relative abundance of nickel were associated with genetic differences. This is most easily achieved when there are either a small number of alleles, or many alleles in physical linkage, associated with nickel abundance. In comparison, magnesium was more continuously correlated with genetic differences, and thus small or large changes in the abundance of this element were associated with genetic effects of similar magnitude. The more linear fit for magnesium could be achieved through the presence of a larger number of independently segregating alleles, consistent with the identification of 79 alleles associated with magnesium tolerant Arabidopsis phenotypes (Niu et al. 2018). In comparison, it is more likely that there are a smaller number of alleles, or more alleles in physical linkage, associated with nickel abundance.

Studies of magnesium and nickel tolerance have focused on mutations of large effect that differentiate serpentine from nonserpentine populations (Bradshaw 2005, Niu et al. 2018, Celestini et al. 2025). However, given that our analyses are based on total genomic differences between individuals, and are not weighted by any phenotypic measurements, our results suggest that adaptation to soil composition occurs across many loci, potentially of varying effect, that are not fixed within the population. This adaptive genetic architecture is consistent with a model of adaptation from standing genetic variation (e.g. soft selection) and allows multiple allelic combinations to produce similar phenotypes, even with the inclusion of large effect alleles (Jones et al. 2012, Yeaman 2015, Tigano and Friesen 2016, Szukala et al. 2021). This genetic architecture is also more likely when alleles have additive effects and lack pleiotropy, a pattern that was identified across 17 plant species experiencing repeated selection (Nocchi et al. 2024). The extreme fine-scale resolution of our study thus suggests that environmental heterogeneity over small spatial scales could play an important role in the development of adaptive genetic architecture, particularly for serpentine plant populations. To our knowledge, associations between intraspecific variation in serpentine genotypes and environmental variation has received little attention compared to studies of broadscale serpentine adaptation. While the importance of this relationship between genetic composition and the local environment in *C. tiburonensis* remains uncertain, our results suggest it plays an important role in the conservation of this, and other, threatened species.

Further genomic analyses will be required to confirm that the patterns observed between soil and genetic composition are adaptive and not merely correlative. Mapping the genetic basis of soil adaptation could confirm the associations depicted here and potentially determine whether alleles are under balancing or directional selection. Additionally, describing transcriptomic variation in *C. tiburonensis* could be useful for pinpointing how genetic composition contributes to fitness across the complex soil environment, particularly if genetic differences result in regulatory changes rather than mutations in protein coding regions. These future analyses would be an essential second step for understanding how soil heterogeneity structures genetic variation in this species. Perhaps more importantly however, it could provide a framework for how varying selection across small spatial scales preserves functional genetic variation broadly.

### Genetic Diversity and Conservation Implications

Differences in collection, sequencing, and analysis make metrics of diversity difficult to accurately compare between species and across studies. However, comparisons with congeners using similar methods to those employed here indicate that *C. tiburonensis* has relatively low overall genetic diversity, consistent with a small but stable species range and population size. For example, *C. tiburonensis* has a larger inbreeding coefficient of 0.205 than the more widespread *C. venustus*, with an F_is_ of 0.107 (Herández et al. 2022). This is unsurprising given the much larger population sizes and range of C. venustus. However, major differences in the scale of sampling (*C. venustus* samples were collected at much greater inter-sample distances) could also cause this pattern and complicate any genomic comparisons. A comparison of nucleotide diversity (*pi*) calculated using all sequenced and assembled sites (including invariant sites) is more appropriate for comparisons across species. Yet this metric is also susceptible to differences in sequencing and analysis methods. In a study using the same sequencing technology (RADseq) and analysis software (STACKS) to calculate all sites *pi*, Rota et al. (2023) studied three range limited species endemic to the Dolomites and Carnic Prealps. They find lower values of *pi* in the two lower elevation species that both experienced range contraction during the last glacial maximum, while the highest elevation species, which also had the most stable population size through time, had higher pi values (0.00137 - 0.00199), very similar to those of *C. tiburonensis* (0.0012 - 0.0018), despite *C. tiburonensis*’ more restricted range and global census population size. These comparisons indicate that *C. tiburonensis* does not have particularly low diversity, that might be maintained by high connectivity (no population structure) and long, overlapping generations that enable the reintroduction and spread of rare alleles.

The simple fact that *C. tiburonensis* is confined to a single hilltop makes it vulnerable to extinction. Translocation and cross pollination are common conservation strategies used to maintain stable population sizes and preserve genetic diversity, but can disrupt genetic structure and local adaptation. However, we found that *C. tiburonensis* exists as a single, large, panmictic population with no structure, meaning that there does not appear to be any risk in utilizing these conservation measures should the need arise in the future.

Additionally, we found no evidence that low genetic diversity contributes to the risk of extinction. However, climate change poses a notable and existential threat to this species, because *C. tiburonensis* is restricted to patchily distributed serpentine soils. Migrating to higher elevations or latitudes to avoid a warming and increasingly arid climate is not possible, so it must adapt in situ or risk extinction. The frequency, severity and duration of droughts in the western US is expected to increase as the climate continues to change (Seager et al. 2007). Fortunately, the relatively high genetic diversity suggests that *C. tiburonensis* has the capacity to adapt to changing environmental conditions.

## Conclusions

Environmental heterogeneity is a common determinant of an organism’s genetic composition (Smith 1997, Parisod and Christin 2008). In plants, this has primarily been shown in response to climatic differences, often over large spatial scales (Gugger et al. 2019). However, local environmental differences can also structure genetic variation at smaller spatial scales in organisms that lack population genetic structure. This phenomenon has been observed most often in marine systems, where large genetic neighborhoods can far exceed the spatial scope of environmental variation (Hoey and Pinsky 2017). Our results suggest that soil heterogeneity, which can exist at very small spatial scales, might drive similar patterns in terrestrial plant populations. Future work using comparative transcriptomics, genome scans, or translocation experiments will be necessary to confirm that selection drives the patterns described by this study.

Nonetheless, the work presented here suggests that variation in soil composition structures plant genetic differences at small spatial scales and could be important for the maintenance of genetic diversity. Clarifying the mechanistic role of selection, in this and other systems, will improve our understanding of the relationship between edaphic heterogeneity and adaptation and allow for the improved management of natural plant populations.

## Author Contributions

SS conceived of the study; SS and JH conducted the field work and collected the data; JB and JH performed the analyses; JB, JH, and SS drafted the manuscript; JB, JH, and SS revised and edited the manuscript.

## Acknowledgements

We thank Jeff Ross Ibara and Katrina Dlugosh for useful feedback on the preliminary results, Geneva Lee for assistance with the extraction and preparation of DNA. This work was supported by Marin County Parks.

